# Metabolite sequestration enables rapid recovery from fatty acid depletion in *Escherichia coli*

**DOI:** 10.1101/590943

**Authors:** Christopher J. Hartline, Ahmad A. Mannan, Di Liu, Fuzhong Zhang, Diego A. Oyarzún

## Abstract

Microbes adapt their metabolism to take advantage of nutrients in their environment. Such adaptations control specific metabolic pathways to match energetic demands with nutrient availability. Upon depletion of nutrients, rapid pathway recovery is key to release cellular resources required for survival in the new nutritional condition. Yet little is known about the regulatory strategies that microbes employ to accelerate pathway recovery in response to nutrient depletion. Using the fatty acid catabolic pathway in *Escherichia coli*, here we show that fast recovery can be achieved by rapid release of a transcriptional regulator from a metabolite-sequestered complex. With a combination of mathematical modelling and experiments, we show that recovery dynamics depend critically on the rate of metabolite consumption and the exposure time to nutrient. We constructed strains with re-wired transcriptional regulatory architectures that highlight the metabolic benefits of negative autoregulation over constitutive and positive autoregulation. Our results have wide-ranging implications for our understanding of metabolic adaptations, as well as guiding the design of gene circuitry for synthetic biology and metabolic engineering.

## Introduction

Bacteria constantly adapt to changing environments by coordinating multiple levels of their intracellular machinery. Metabolic regulation provides a control layer that adapts metabolic activity to nutritional conditions. Such regulation relies on a complex interplay between gene expression and metabolic pathways [1]. In the case of metabolic pathways, genes for nutrient uptake and consumption need to be upregulated when the specific nutrient is available in the environment. Failure to quickly increase pathway capacity may result in missed metabolic resource opportunity and a potential cost on fitness [2] and population survival [3]–[5]. Conversely, upon nutrient depletion, expression of specific metabolic enzymes can become wasteful and lead to a suboptimal use of biosynthetic resources [6], [7].

Metabolite-responsive transcription factors are a widespread regulatory mechanism in microbes. Upon sensing nutrient availability, they trigger changes in enzyme expression and metabolic flux [8]. This strategy has been shown to control the dynamics of pathway upregulation in various ways [9]–[11]. For example, negative autoregulation of transcription factors can speed the response time of gene expression [12] and feedback circuits based on metabolite-responsive transcription factors have been demonstrated to accelerate metabolite responses [13]. While much of the literature has focused on the control of activation dynamics upon nutrient induction [14]–[16], little is known on how these regulatory mechanisms shape pathway recovery after depletion of nutrients.

Here we study a common regulatory architecture found in over a dozen bacterial nutrient uptake systems [17] (Figure 1A and Supplementary Table S1). When a nutrient is absent from the environment, a metabolite-responsive transcription factor (MRTF) represses the expression of uptake and catabolic enzymes. When the nutrient is present, the nutrient is internalized and sequesters the transcription factor via reversible binding, thus preventing gene repression. This causes an upregulation of metabolic enzyme genes and an increase in the rate of nutrient import and utilization. A common feature of these control systems is the presence of negative autoregulation of the transcription factor (Supplementary Table S1). After nutrient depletion, the MRTF must recover its repressive activity on the catabolic pathway genes to rapidly shut down pathway activity, yet it is unclear what components of the regulatory system help to accelerate the recovery dynamics.

**Figure 1.**
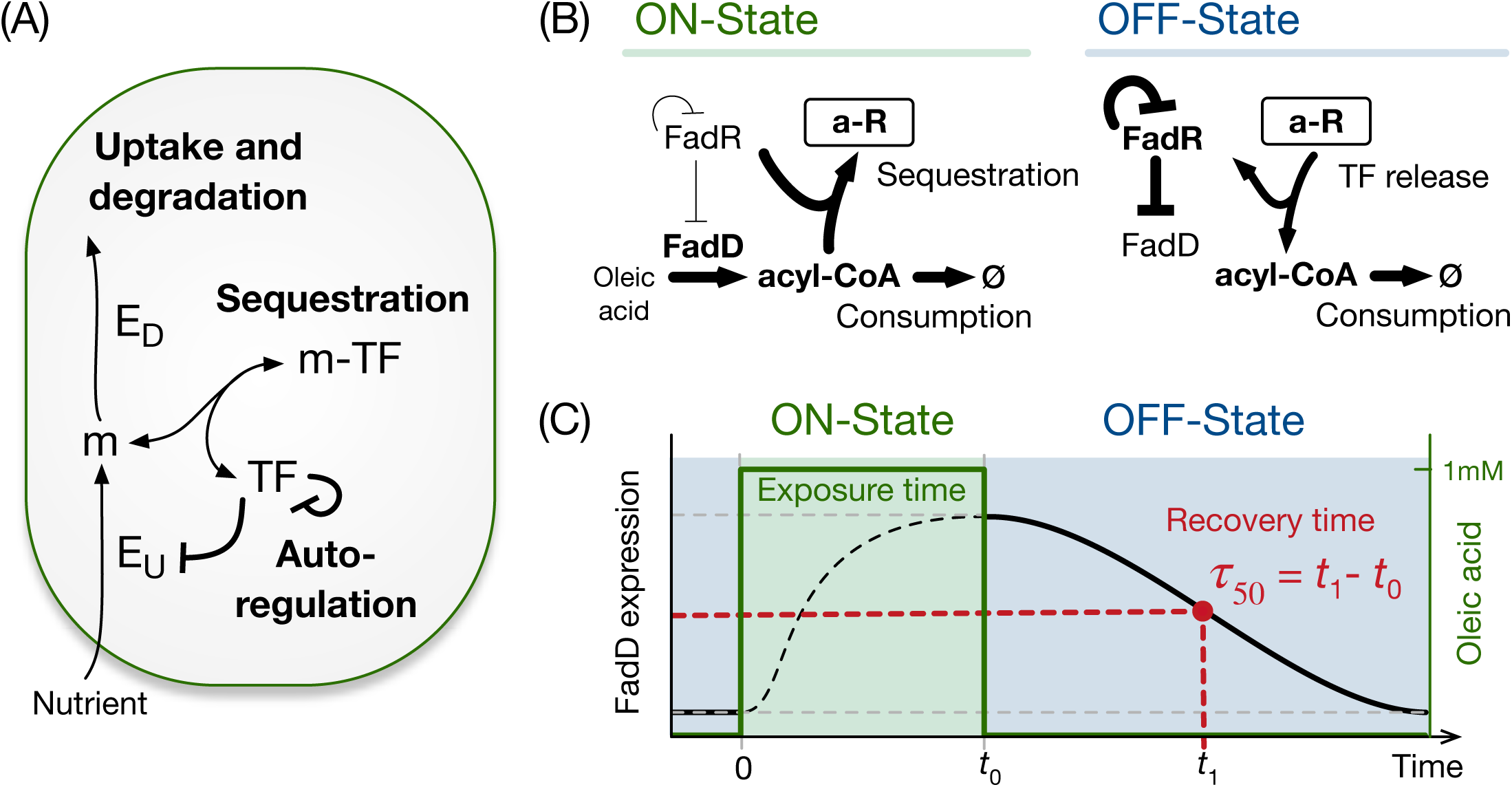
General architecture of a bacterial nutrient uptake system. **(A)** Regulation of nutrient uptake by a metabolite-responsive transcription factor, a ubiquitously observed control system in bacteria (Supplementary Table S1). **(B)** We use the *Escherichia coli* fatty acid uptake as a model system. The ON-state is defined by induction at a constant level of oleic acid, which is imported as acyl-CoA by uptake enzyme FadD. Acyl-CoA sequesters the transcription factor FadR, which de-represses expression of the uptake enzyme. The OFF-state is defined by the wash out of oleic acid after some time *t*_0_ in the ON-state. Release of sequestered FadR recovers its repression on FadD synthesis. FadR is also subject to negative autoregulation. **(C)** Schematic of the experiments and simulations in this work, with defined exposure time to oleic acid (green area), and recovery time of FadD levels in the OFF-state (*τ*_50_) defined as the time to reach to half way between maximum and minimum concentrations.

Using the *Escherichia coli* fatty acid catabolic pathway as our model system, we took a theoretical-experimental approach to study its recovery dynamics in response to a nutrient shift from an ON-state to an OFF-state. As illustrated in Figure 1B, these two states are defined as an environment with and without the presence of oleic acid as carbon source, respectively. In the ON-state, oleic acid is imported as fatty acyl-CoA which binds to the transcription factor FadR and sequesters it into a complex. This acyl-CoA sequestration releases FadR from its cognate DNA elements [18], which relieves the repression of the uptake gene *fadD* and thus accelerates the import of oleic acid. We found that upon depletion of oleic acid, repression by FadR is recovered via its rapid release from the sequestered complex, which in turn is driven by consumption of acyl-CoA. We further found that the architecture of FadR autoregulation affects the maintenance of a sequestered pool of FadR. In particular, negative autoregulation enables a large sequestered TF pool during the ON-state and, at the same time, a reduced biosynthetic cost in the OFF-state. Our results shed light on the regulatory mechanisms that allow cells to rapidly adapt to environmental shifts and provide insights for the design of gene circuits in synthetic biology and metabolic engineering applications, particularly where strain performance is sensitive to nutrient fluctuations and inhomogeneities typical of large-scale fermentations.

## Results

### Recovery dynamics in the fatty acid uptake system

To study the recovery dynamics of fatty acid uptake, we built a kinetic model based on four core components of the regulatory system: FadD (*D*), free FadR (*R*), acyl-CoA (*A*) and sequestered FadR (*aR*). The model represents cells growing at a fixed growth rate with oleic acid at a fixed concentration in the media. We simulated the recovery dynamics by mimicking the three stages in our experimental setup: preculture without oleic acid, response to induction in the ON-state, and recovery in the OFF-state. During preculture we ran the model to steady state in the absence of oleic acid, then initiated simulations of the ON-state from the steady state achieved in preculture, with a fixed concentration of oleic acid for a defined exposure time. The concentrations achieved at the end of the ON-state were used as initial conditions for the OFF-state, which was simulated without oleic acid until the system recovers to the steady state in preculture (Figure 1C).

We defined two metrics to quantify the recovery dynamics after the switch from ON-to OFF-state (Figure 1C). First, we define the recovery time (*τ*_50_) as the time taken for FadD to decrease to half-way between its maximum and minimum steady state value after nutrient depletion (Figure 1C). Second, we defined the metric *η* as the proportion of free FadR released from the sequestered complex after one doubling time:

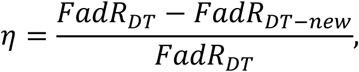

where FadR_DT_ and FadR_DT-new_ are the concentrations of free FadR and newly expressed FadR in the OFF-state after one doubling time (DT). This definition allows us to quantify the contribution of free FadR released from the sequestered pool to the recovery dynamics.

Since pathway recovery depends on the system state at the time of the ON to OFF switch, we used the kinetic model to study the relation between the initial conditions at the time of the switch and the recovery dynamics. To this end, we studied the impact of exposure time to oleic acid during the ON-state, as well as the amount of acyl-CoA-consuming enzyme. We simulated the OFF-state dynamics for 2500 combinations of 50 acyl-CoA-consuming enzyme concentrations and 50 exposure times, and calculated *τ*_50_ and *η* for each. Simulation results of the OFF-state dynamics (Figure 2A) suggest that the recovery time (*τ*_50_) decreases with increasing concentrations of consuming enzyme, while the amount of released FadR (*η*) increases with both the consuming enzyme and the exposure time. Further simulations suggest that when exposure time increases, the pool of acyl-CoA accumulates further, with a rise time between 8.5 to 10 hours, for levels of consuming enzyme between 100 μM to 6μM (Figure S2B). This larger pool takes a longer time to be consumed in the OFF-state (Figure S2A) and so delays the release of FadR from the complex. This results in a longer recovery time (details in Supplementary Methods S1 and Figure S2). Model simulations also reveal a strong inverse relation between *τ*_50_ and *η* (Figure 2B), indicating that the release of FadR from sequestration by acyl-CoA provides a mechanism for cells to achieve rapid recovery during nutrient depletion. Further, the sensitivity of this inverse relation increases when cells are exposed to a longer ON-state. Simulations show that longer cell exposure times to oleic acid increase the pool of sequestered FadR (Supplementary Figure S2). Consequently, in the OFF-state, more FadR can be released from sequestration compared to new FadR synthesis, thus increasing the sensitivity of *τ*_50_ changes in the amount of released FadR (*η*).

**Figure 2.**
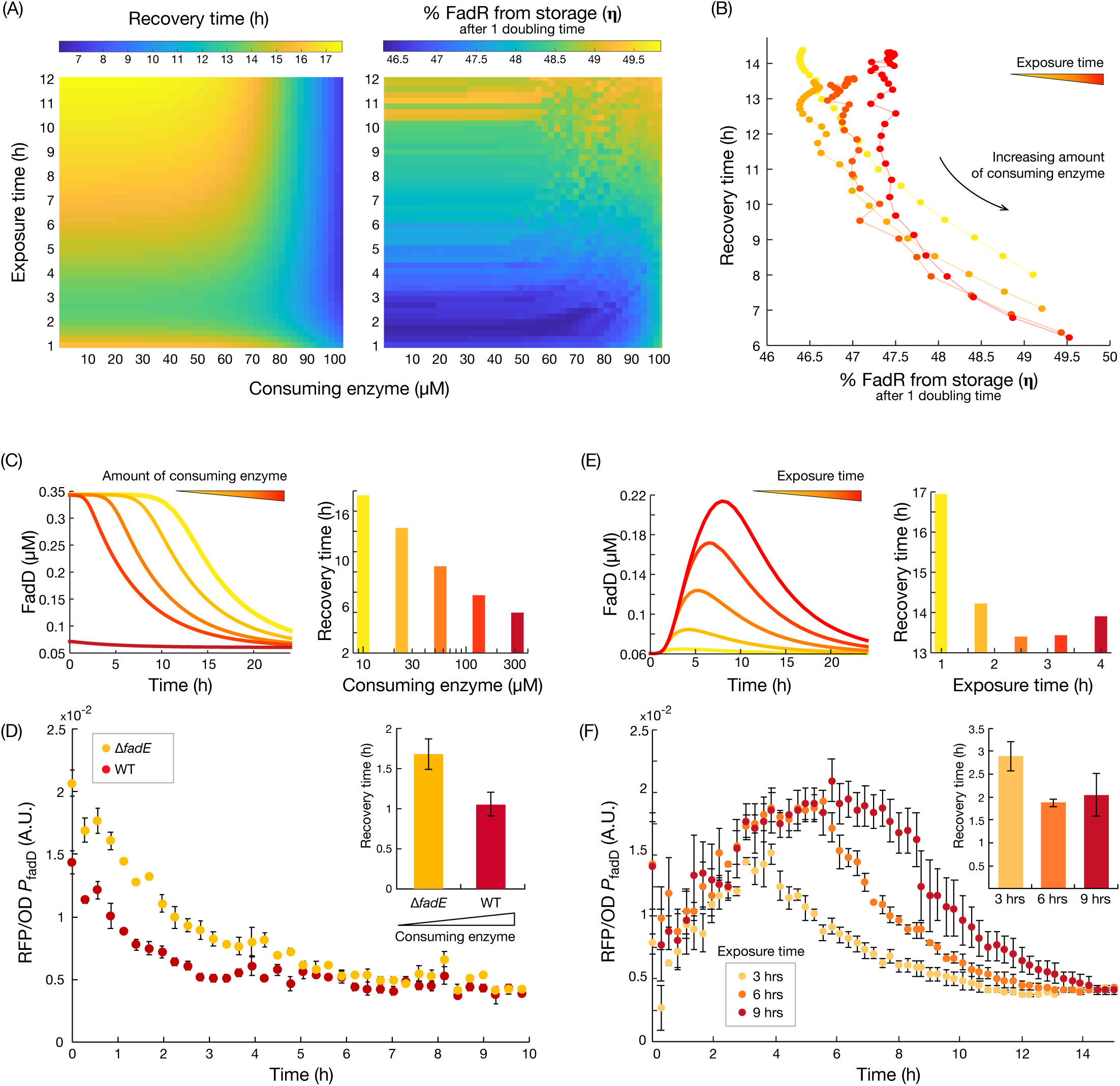
Nutrient exposure time and speed of metabolite consumption in the OFF-state shape the recovery time. **(A)** Predicted recovery time (*τ*_50_) and proportion of free FadR released from sequestration after one doubling time (*η*) for variations in the amount of consuming enzyme and nutrient exposure time. **(B)** Inverse relation between the proportion of released FadR (*η*) and predicted recovery time. **(C)** Simulated time course of FadD concentration during OFF-state and predicted recovery times for increasing concentration of acyl-CoA consuming enzyme. **(D)** Measured time course of *fadD* expression when switching from ON- to OFF-state for strains with low (Δ*fadE*-reporter) and high (WT-reporter) concentration of acyl-CoA consuming enzyme. Strains were switched from M9G+1mM oleic acid to M9G media at time zero. Error bars represent standard error of the mean (SEM) from biological triplicates (n=3). Recovery times were calculated from exponential fits to each of the triplicate time course data (inset). Error bars represent SEM from biological triplicates (n=3). (E) Time course simulations of FadD induction and recovery dynamics, and predicted recovery times, for increasing exposure times. (F) Measured time course of *fadD* expression from the WT-reporter strain grown for 3, 6 and 9 hours of exposure to oleic acid (M9G+1mM oleic acid), and then switched to OFF-state (M9G). Error bars represent SEM from biological triplicates (n=3). Recovery times were again calculated from exponential fits, with error bars SEM from triplicates (n=3).

To verify the model predictions, we sought to experimentally perturb the fraction of released FadR (*η*) through two complementary strategies: i) by engineering strains with different amounts of acyl-CoA consuming enzymes, and ii) by manipulating the exposure time to oleic acid. We first constructed a reporter strain with a decreased consumption rate of acyl-CoA, strain ΔfadE-reporter (see Supplementary Table S3B), where we deleted the *fadE* gene encoding the second step of the fatty acid β-oxidation pathway. This prevents metabolization of acyl-CoA by β-oxidation, and leaves membrane incorporation (catalyzed by enzyme PlsB) as the only pathway for acyl-CoA consumption. We measured *fadD* expression dynamics after switching the strains from the ON-state (M9G + 1mM oleic acid media) to OFF-state (M9G media) using a red fluorescent protein (RFP) reporter fused downstream of the *fadD* promoter. The *fadE* knockout strain displayed a slower recovery than the wild type, with ∼60% increase in recovery time (Figure 2D), confirming our theoretical prediction in Figure 2C. The measured increase in recovery time entails an increased expenditure of biosynthetic resources to import a metabolite that is no longer present in the environment.

Next, we measured the *fadD* recovery dynamics after switching the cultures from growth with 3, 6, and 9 hours of exposure time in the ON-state. As predicted from the model in (Figure 2E), measured recovery time decreased for an increase in exposure time (Figure 2F). However, we observe that recovery time is not decreased further beyond 6 hours of exposure to oleic acid. We speculate that faster recovery is counteracted by the delay of having to consume a higher level of accumulated acyl-CoA, or because the maximum level of sequestered FadR may already have been achieved at 6 hours.

### Impact of autoregulatory architecture on recovery dynamics

Among the uptake systems in *E. coli* with the architecture of Figure 1A, we found that the majority have a transcriptional regulator that represses its own expression, few have constitutive expression of the regulator, and none display positive autoregulation (see Supplementary Table S1). To better understand the salient features of each regulatory architecture and how they affect recovery dynamics, we built variants of our kinetic model with FadR under constitutive expression and positive or negative autoregulation (details in Methods). Simulations of the recovery dynamics in the OFF-state for varying exposure times in the ON-state suggest that these architectures behave similarly for short exposure times (< 1 hour), quickly sequestering all the free FadR (Figure 3A, top). For longer exposure times (>1 hour), model simulations suggest important differences in the dynamics of sequestered FadR among the various modes of autoregulation. Negative autoregulation shows an accumulation of sequestered FadR, while positive autoregulation leads to an overall depletion of sequestered FadR. Constitutive expression causes the total level of sequestered FadR to be maintained at a constant level (Figure 3A).

**Figure 3.**
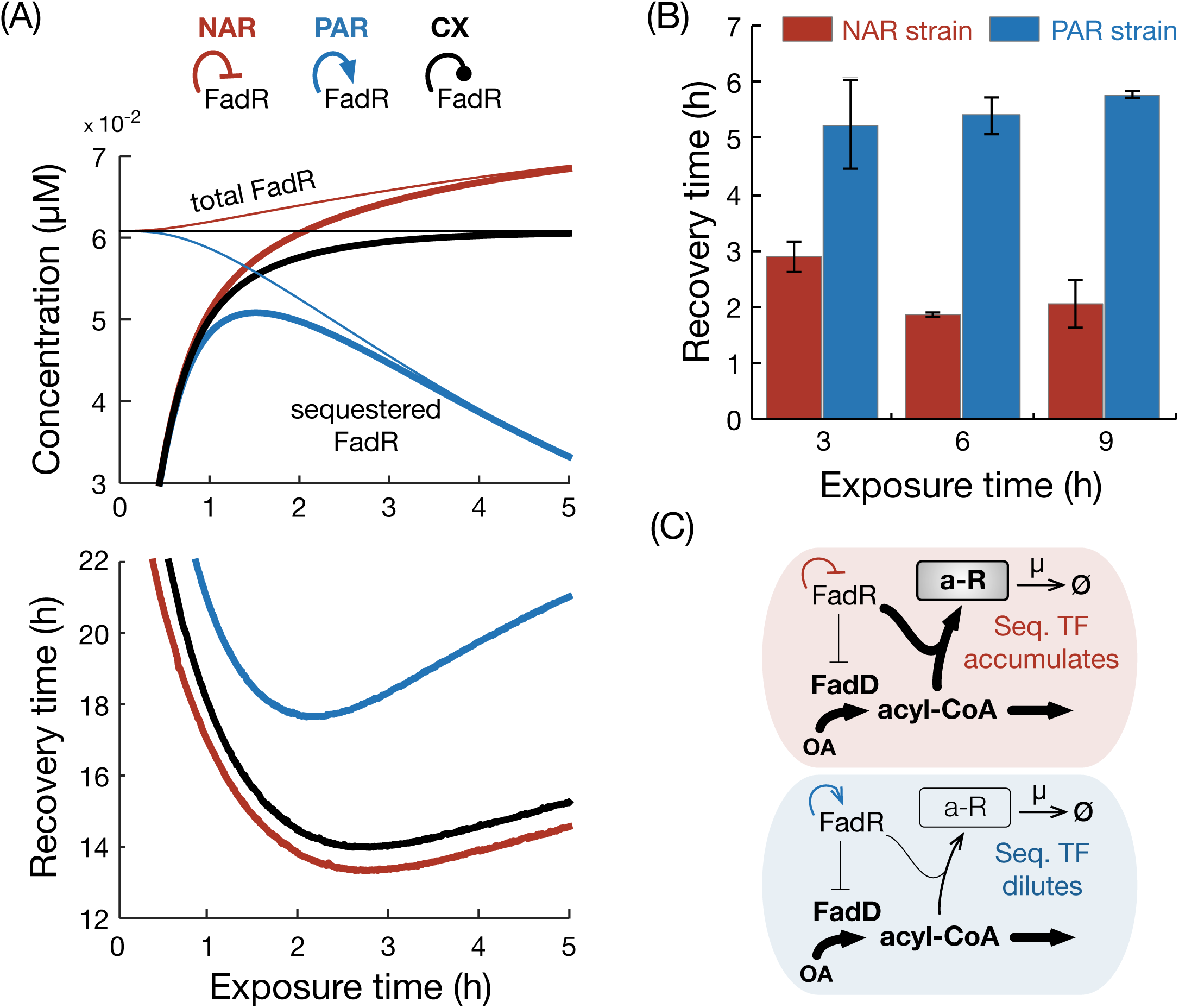
Impact of regulatory architecture on the recovery time after nutrient depletion. **(A)** Top: Simulated steady state concentrations of sequestered (thick line) and total FadR (thin line) for varying times spent in the ON-state for three regulatory architectures of FadR; constitutive expression (black line) is represented by a blunt line. Bottom: Predicted recovery times for each architecture. **(B)** Measured recovery times in the WT (WT-reporter) and positively autoregulated strain (PA-reporter, Supplementary Table S3B and Table S4) for 3, 6 and 9 hours of exposure in ON-state. Recovery times were calculated from exponential fits to each of the triplicate time course data (see Supplementary Methods S1 and Figure S3) and error bars represent SEM of the calculated values (n=3). **(C)** Schematics illustrating how negative and positive autoregulation affect the build-up of sequestered FadR in the ON-state.

To elucidate whether these predicted trends are a consequence of the model parameters or inherently determined by the autoregulatory architecture, we analyzed the model and found relations for the change in steady state concentrations of total FadR (Δ*R*_*T*_) in each autoregulatory architecture (details of derivation in Supplementary Methods S1):

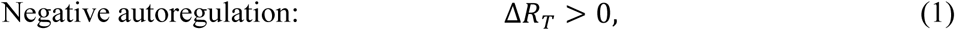

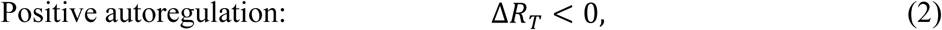

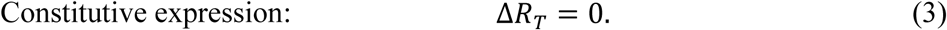

Equations (1), (2) and (3) are valid for any combination of positive parameters, and therefore the long-term trends observed in Figure 3A are structural properties of the model.

To determine the effect of the three regulatory architectures on the recovery time, we simulated the recovery dynamics of each architecture for varying exposure times and calculated the recovery time (Figure 3A, bottom). We observe that the overall relation between recovery time and exposure time is similar across the three architectures (Figure 3A, bottom inset). However, for positive autoregulation we found recovery to be significantly slower for a wide range of exposure times. To test this prediction, we engineered an *E. coli* strain with positively autoregulated FadR expression by replacing the native *fadR* promoter with one that activated by FadR (P_*fadRpo*_, see Supplementary Table S4), and a P_fadD_ reporter plasmid. The positively autoregulated reporter strain (PA-reporter, Supplementary Table S3B and Table S4) was grown in the ON-state (M9G media + 1mM oleic acid) and then rapidly switched to the OFF-state (M9G media) after 3, 6 and 9 hours. We measured the *fadD* expression dynamics (see time course dynamics in Supplementary Figure S3B), and calculated the respective recovery times (Figure 3B). Consistent with the trend predicted from the model, recovery times for the positively autoregulated strain increased with the exposure time of oleic acid in the ON-state (Figure 3C).

### Negative autoregulation provides a resource-saving recovery strategy

The results in Equations (1) and (3) suggest that constitutive expression and negative autoregulation can both maintain high amounts of sequestered FadR for long exposure times to oleic acid. Our earlier results showed that longer exposure times lead to larger pool of sequestered FadR (Supplementary Figure S2D), which enables a faster recovery time (Figure 2E, F). We thus asked which system parameters influence the steady-state pool size of sequestered FadR in these two architectures. We found that for high concentrations of oleic acid, the steady-state concentration of sequestered FadR in the ON-state are given by (details in Supplementary Methods S1):

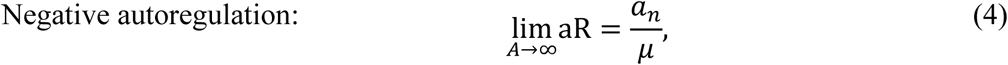

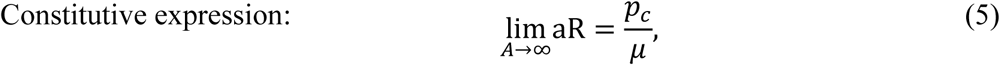

where A and aR are the steady state concentrations of acyl-coA and sequestered FadR, respectively. These results suggest that at high oleic acid concentrations, the amount of sequestered FadR scales linearly with the strength of its own promoter. Simulations of both architectures in the ON-state induced with high concentration of oleic acid (1mM) and varying promoter strengths, we found that increasing promoter strength both increases the amount of sequestered FadR in the ON-state and decreases the recovery time (Figure 4A).

**Figure 4.**
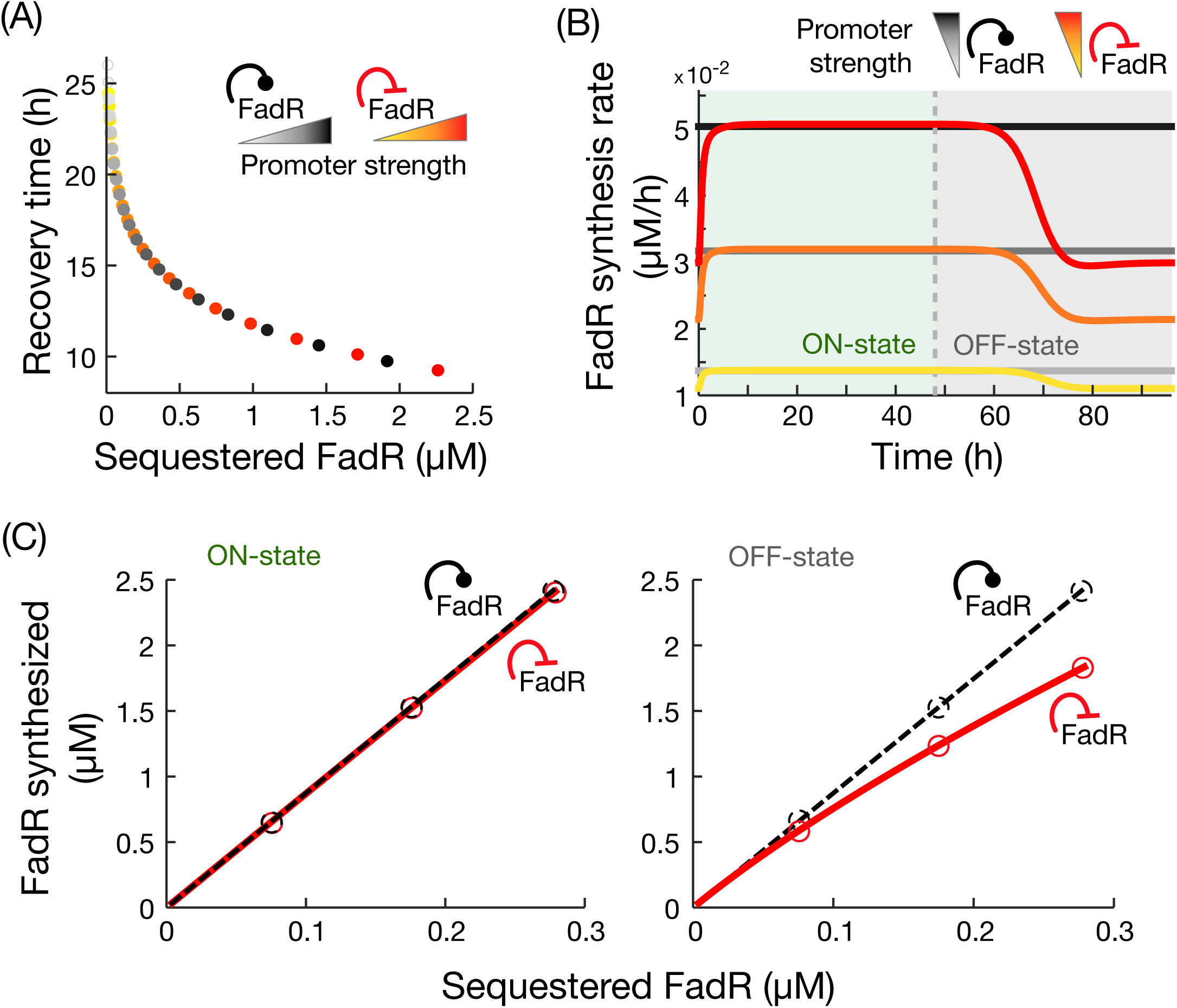
Comparison of recovery dynamics in constitutive expression and negative autoregulation. **(A)** Simulated recovery times for variations in the strength of FadR’s own promoter, with both architectures achieving the same recovery times. **(B)** Time course simulations of FadR synthesis rates for 48 hours in the ON-state (1mM oleic acid) and OFF-state, for increasing promoter strengths; yellow curve represents response with the fitted promoter strength value (Supplementary Table S2B). To ensure fair comparison, promoter strengths were chosen to achieve the same recovery time in the two architectures. **(C)** Cost of FadR synthesis for increasing concentrations of sequestered FadR, modified by changes to *fadR* promoter strength. Circles correspond to costs associated to simulations shown in (B). Details of simulations in Methods.

The results in Figure 4A also suggest that through tuning of *fadR* promoter strength, in principle both constitutive expression and negative autoregulation can produce the same recovery time. We thus sought to identify potential benefits of one architecture over the other in terms of the recovery dynamics in the OFF-state. Since production of FadR entails a biosynthetic cost, we compared both regulatory architectures in terms of the cost of FadR synthesis. From time-course simulations of FadR synthesis rates in the ON- and OFF-states (Figure 4B), we computed the total amount of synthesized FadR for increasing *fadR* promoter strengths by integrating the area under the curves (Figure 4C). Our results show that both architectures require identical biosynthetic costs for FadR in the ON-state, but negative autoregulation leads to a reduced biosynthetic cost for FadR in the OFF-state as compared to constitutive expression (Figure 4C).

## Discussion

In this paper we combined mathematical modelling and experiments to study metabolic pathway recovery upon depletion of an external nutrient. Changes in nutrient conditions trigger transcriptional programs that adapt cell physiology [19] to meet the cellular energy budget [20]. We chose the regulation of fatty acid uptake in *E. coli* as our model system, as it is representative of a widely conserved transcriptional program for controlling the uptake of nutrients in bacteria (Supplementary Table S1). We show that fast recovery after nutrient depletion can be achieved by rapid release of a transcriptional regulator from a metabolite-sequestered complex. In particular, a sizable contribution of FadR rapidly made available after oleic acid depletion came from its release from its sequestered complex form (aR), as opposed to new synthesis. The rapid availability of FadR quickly recovers its inhibition on the *fad* regulon, and so shortens the recovery time. Furthermore, our model simulations and experiments have demonstrated that increasing the amount of FadR stored in complex form during nutrient exposure and fast consumption of acyl-CoA (the sequestering metabolite) facilitate a speedy recovery in the OFF-state.

Our model simulations show that pathway recovery is delayed by high intracellular acyl-CoA concentrations, which slow the release of free FadR from stored complex until those high concentrations are reduced. This delay occurs because FadR is only able to sense the intracellular metabolite concentrations, which can remain high even when extracellular metabolite concentrations are low. During this delay, wasteful expression of the uptake pathway continues despite the absence of oleic acid in the environment. Previous research has shown that upon nutrient induction, metabolite dynamics tend to lag behind slow upregulation of metabolic enzymes [13]. In contrast, here we find that after inducer depletion, the recovery of metabolic enzymes back to their downregulated state lags behind the metabolite dynamics. This has important implications for designing synthetic control circuits which utilize non-metabolizable inducers such as IPTG or TMG. Without consumption of the inducer, post-induction recovery response will be slow and may cause a dramatic drain of cellular resources. Our simulations of the relation between sequestered FadR and recovery time suggest that this inherent lag can be compensated for by storing and releasing higher amounts of TF, which highlights the benefits of maintaining a sequestered pool of FadR.

Further mathematical analyses revealed principles that explain how autoregulation shapes the recovery time. We found that systems with only negative autoregulation and constitutive expression can maintain the pool of sequestered FadR needed for a rapid recovery. In contrast, we found that positive autoregulation loses this storage over time, resulting in a reduced availability of FadR after nutrient depletion and slower recovery times. We additionally found that negative autoregulation of the transcription factor reduces the total biosynthetic cost of for FadR in a full ON-OFF-state cycle as compared to using constitutive expression. This occurs because both systems need to maintain the same level of sequestered FadR in the ON-state in order to achieve the same recovery time, but only negative autoregulation allows FadR synthesis to be down-regulated in the OFF-state. Thus, negative autoregulation provides a resource-saving strategy for controlling the recovery dynamics compared to constitutive expression. We found that the transcriptional regulators in 13 out of 18 nutrient uptake systems (see Supplementary Table S1) have negative autoregulation, suggesting an evolutionary pressure for a resource-saving control strategy. Past studies in the literature have found that expression under negative autoregulation can decrease response times in gene expression [12], linearize dose-response in responsive systems [21], and even speed up metabolic dynamics [13]. In addition to these properties, we find that negative autoregulation enables a rapid and more resource-saving metabolic recovery to nutrient depletion.

Recent efforts in synthetic biology focus on engineering gene control circuits to manipulate microbial metabolism [22]–[24]. One key goal of such control systems is to rapidly turn off metabolic pathways in response to metabolic signals [25]–[27]. Our results provide core design principles for engineered metabolic systems with tunable response to nutrient depletions, which could be used as a pathway control tool in bioreactors. Our experiments and simulations reveal that the recovery time can be simply tuned through well-established promoter engineering techniques [28]–[30]. Further, we identify regulatory architectures with differing dynamic responses to nutrient depletion provides further avenues for tuning system response to the highly dynamic and heterogeneous environments typical of large-scale fermenters. These design rules can be readily applied to mitigate against deleterious nutrient fluctuations found in metabolic engineering applications.

## Materials and Methods

### Materials

Phusion DNA polymerase, T4 DNA ligase, restriction enzymes, and Teknova 5x M9 minimal salts were purchased from Thermo Fisher Scientific (Waltham, MA, USA). Gel purification and plasmid miniprep kits were purchased from iNtRON Biotechnology (Lynnwood, WA, USA.). Oligonucleotides were synthesized by Integrated DNA Technologies (Coralville, IA, USA). All other reagents were purchased from Sigma-Aldrich (St. Louis, MO, USA.)

### Plasmids, strains, and genome modifications

A list of plasmids used along with promoter sequences in this study is provided in Supplementary Methods S1, Table S3 and Table S4. *E. coli* DH10β was used for plasmid construction. The plasmid pSfadDk-rfp was constructed by cloning the *fadD* promoter (500 bp upstream of its translation start site) into the 5’ of a *rfp* gene in a BglBrick vector, pBbSk-rfp [31] using Golden Gate DNA Assembly [32]. The positively autoregulated *fadR* strain was engineered by replacing *fadR*’s native promoter with a FadR-activated promoter P_*fadRpo*_ via CRISPR-Cas9 genome editing [33]. Detailed engineering methods and the characterization of the P_*fadRpo*_ promoter are described in Supplementary Methods S1.

Three reporter strains were created to measure expression dynamics from the *fadD* promoter. These strains were created by transforming plasmid pSfadDk-rfp into either the wild-type DH1 strain, DH1(ΔfadE), or an engineered strain with positively autoregulated *fadR*, resulting in WT-reporter, ΔfadE-reporter, and PA-reporter, respectively.

### Media conditions

All strains were grown from single colonies and cultivated overnight in Luria-Bertani (LB) media before experiments. For OFF-State culture conditions, cells were grown in M9 minimal media [34] supplemented with 1% glycerol and 0.5% Tergitol Solution Type NP-40 (M9G). For ON-state culture conditions, cells were grown in M9G + 1 mM oleic acid (M9G+OA). All cultures were supplemented with appropriate antibiotic selection (50 mg/L Kanamycin, 100 mg/L Ampicillin).

### Assays of fadD expression dynamics

To measure the recovery dynamics, reporter strains were grown in 3 mL M9G+OA for 24-48 hours at exponential growing state. To rapidly switch nutrient, cells were centrifuged (5500 rcf, 2 minutes) and washed twice in M9G. Cultures were then diluted in M9G medium to OD_600_ = 0.08 and transferred to a Falcon 96-Well Imaging Microplate (Corning, NY, USA). The microplate was then incubated in an Infinite F200PRO plate reader (TECAN, Männedorf, Switzerland) at 37°C with constant shaking. To maintain exponential growth during measurement, cultures were diluted by a factor of 5 for three times during incubation. Kinetic measurements of cell density (absorbance at 600 nm) and RFP fluorescence (excitation: 584 ± 9 nm, emission: 620 ± 20 nm) were taken every 900 seconds until all diluted cultures reached stationary phase. Fluorescence from water in the same 96-well plate was used as the background and was subtracted from all fluorescence measurements. The background-corrected fluorescence was later normalized by cell density. To calculate the recovery time, the average of three biological replicates were fitted to an exponential curve:

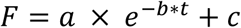

where *F* is the background-corrected, cell-density-normalized fluorescence. The recovery time was calculated as *τ*_50_=log(2)/*b*.

For switches after defined times in the ON-state, cultures were first grown in exponential growth phase for 24-28 hours in M9G. Samples from these cultures were then centrifuged (5500 rcf, 2 minutes) and suspended in M9G+OA with an initial OD_600_ of 0.08 and cultivated in 96-well plates for various amount of time as indicated.

### Kinetic model of fatty acid uptake

To study the dynamic response to oleic acid exposure (ON-state) and its recovery (OFF-state) (Figure 1C), we built a kinetic model of the fatty acid uptake system. We define the model as a system of ODEs describing the rate of change of each species:

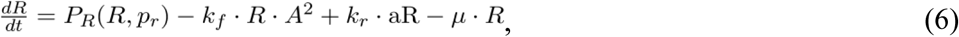

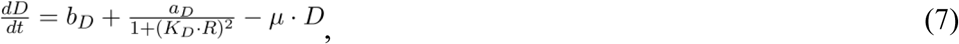

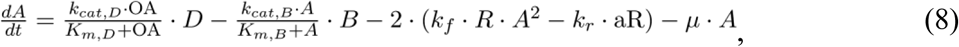

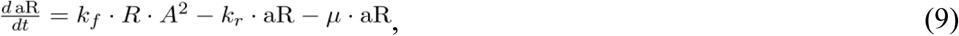

where *R, D, A* and aR represent the concentrations of transcription factor FadR, uptake enzyme FadD, internalized fatty acid acyl-CoA, and sequestered complex acyl-CoA-FadR, respectively (Figure1B). The reversible sequestering of one FadR dimer by two acyl-CoA molecules (stoichiometry as defined in [35]) is modeled as mass-action kinetics in the term *k*_*f*_ *RA*^2^ − *k*_*r*_aR. The term *P*_*R*_(*R,p*_*r*_) represents the expression and autoregulation of the fadR promoter. To model FadR negative autoregulation for the wild-type strain, we use:

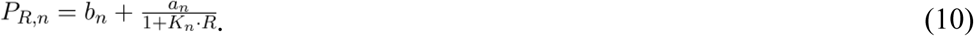

To fit model parameters, we first extended the model to simulate batch culture, and then applied a least-squares fitting of simulations to time course measurements of RFP fluorescence expressed under a fadD promoter, from strain ΔfadE, in various concentrations of oleic acid (see details in Supplementary Methods S1 and Table S2A). Fitting results are illustrated in Supplementary Figure S1 and the fitted parameter values are reported in Supplementary Table S2B. These values were used throughout this study, unless otherwise stated. To understand the impact of model parameters on the recovery time, we performed global parameter sensitivity analysis (details in Supplementary Methods S1 and Figure S4). To model the strains with positive autoregulation and constitutive expression of FadR, we use:

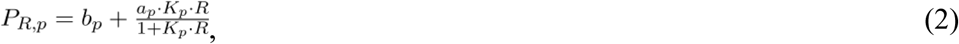

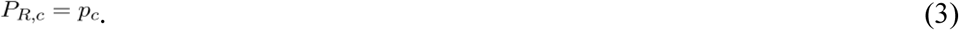

### Model simulations

The model was solved with the MATLAB R2018a ODE solver suite. To simulate the ON-state: simulations were initialized using steady state values achieved from simulations of the preculture (oleic acid, OA = 0 µM), and setting a constant oleic acid concentration of 1000 µM. Simulations were then run for a defined exposure time. To simulate the OFF-state: the system was initialized from the state achieved at the end of the ON-state, and oleic acid set to 0 µM. Simulations were then run to steady state, and recovery times were calculated as the time from the start of the OFF-state till FadD reached half-way between its initial value and minimum steady state value. To calculate the cost of FadR synthesis in the ON and OFF-states (Figure 4C), we integrated simulations of FadR synthesis rate over 48hrs in each state.

In Figure 3, for fair comparison model parameters are set such that the steady state concentration of FadR is the same for all three architectures prior to switching to the ON-state. Likewise, in Figure 4B-C for fair comparison, *fadR* promoter strengths for both architectures were set to achieve same concentration of sequestered FadR in the ON-state (and thus equal recovery times).

## Acknowledgements

This work was funded by the Human Frontier Science Program through a Young Investigator Grant awarded to F.Z. and D.O (grant no. RGY-0076-2015) and by the US National Science Foundation (MCB1453147) to F.Z.

## Author contributions

C.J.H. and F.Z. designed experiments. A.A.M and D.A.O. designed the mathematical analysis. C.J.H. and D.L. engineered strains, carried out experiments, and interpreted the data. A.A.M. developed models, parameter fitting and sensitivity analysis, and model analyses. F.Z. and D.A.O. designed and supervised the study. All authors drafted the manuscript.

## Supplementary Information

**Supplementary Methods S1.** Details on experimental methods, model fitting and mathematical analysis.

**Supplementary Figure S1. (A)** Plot of time series data of measured optical density (OD), in log scale, during growth in media induced with titrations of oleic acid. Values of the fitted growth rate to each data series is given inset, including average growth rate (based on all data series) used to in the model; modelled growth shown in red line, ± SEM in red dashed lines. **(B)** Ensemble of 100 independent fits (gray curves) and the optimal fit (coloured curves) of simulations to time course data (point with errorbars), after converting fluorescence values to concentration. Error bars in data represent SEM from biological triplicates (n=3). Fitting performance in inset.

**Supplementary Figure S2.** Exposure time to oleic acid affects accumulation of acyl-CoA and the recovery time. Coloured curves show how the respective relations are affected by increases in the concentration of consuming enzyme. Inset bar plot in (B) shows rise times of the accumulating acyl-CoA for increases in the consuming enzyme, where rise time is defined as the time taken for acyl-CoA to reach half of its steady state level.

**Supplementary Figure S3.** Characterization and use of PA-reporter strain. **(A)** Dose-response of P_fadRpo_-rfp indicates FadR activated and OA inhibited P_*fadRpo*_ expression. Error bars are SEM for biological replicates (n=3) and blue curve is fit of a hill equation to the mean oleic acid concentrations. **(B)** Time course fluorescence data for the switching experiment of the PA-reporter strain. Cells were induced by 1mM oleic acid at time zero, and grown for three different exposure times, 3, 6, and 9 hours (dashed vertical lines), after which cultures were rapidly switched to fresh media lacking oleic acid (OFF-state). Error bars represent the SEM from biological triplicates.

**Supplementary Figure S4.** Global sensitivity analysis of recovery time to model parameters. Bar plot of the total-order sensitivity indices calculated from global sensitivity analysis (GSA) with eFAST. Sensitivities were calculated from 257 samples per search curve (set of parameters), and this sampling was repeated 7 times to ensure coverage of parameter values. Bars and error-bars show the average and 1 standard deviation of sensitivities over the 7 repeated sampling. eFAST assigns a dummy parameter a small, non-zero sensitivity (last bar). This was exploited to perform a two-tailed *t*-test to calculate whether the sensitivity of each parameter was significantly greater than that of the dummy parameter (asterisk, using a significance *α=0.01*). Parameters are listed in descending order of sensitivity (bottom).

**Supplementary Table S1.** List of metabolite-responsive transcription factors (TF) that control expression of nutrient uptake enzymes in *Escherichia coli*, taken from the EcoCyc database. All these systems follow the schematic as our model system illustrated in Figure 1A.

**Supplementary Table S2. (A)** Parameters of the kinetic model. **(B)** Results of parameter fitting. Optimal parameter values together with search bounds and summary statistics for 100 independent fits. The bounds on growth rate, *μ*, are based on ± two times the SEM from data (shown in Supplementary Figure S2A, inset). Hill coefficients are fixed to *n*_*R*_=1 and *n*_*D*_=2, based on the number of FadR binding sites on the fadR and fadD promoters. Concentration of PlsB is fixed to 0.1369 μM, as taken from Schmidt et al. (2016), Nat. Biotechnol, 34(1).

**Supplementary Table S3. (A)** Plasmids used in this study. **(B)** Strains used in this study.

**Supplementary Table S4.** Sequences of engineered promoter with positively autoregulated *fadR*. **(A)** Native *fadR* promoter sequence, P_*fadR*_. Bold lettering indicates FadR operator site, blue lettering indicates coding sequence. **(B)** Positively autoregulated *fadR* promoter, P_*fadR*po,_ engineered in this work. The underlined sequence is derived from the *fabA* promoter region of *E. coli* DH1 genome. **(C)** Engineered P_*fadR*po_ used to control *rfp* expression.

