## Supplemental Text for "Metabolite sequestration enables rapid recovery from fatty acid depletion in *Escherichia coli*"

#### Table of Contents

**S1** Nutrient uptake systems in *Escherichia coli*
**S2** Kinetic model of fatty acid uptake
**S3** Impact of exposure time to nutrient
**S4** Plasmids and strains used in this study
**S5** Steady state analysis: autoregulation affects FadR levels during induction
**S6** Construction and characterization of strain with positively autoregulated *fadR*
**S7** Steady state analysis: impact of promoter strength on sequestered FadR
**S8** Sensitivity analysis of kinetic model

### S1. Nutrient uptake systems in *Escherichia coli*

**Table S1.** List of metabolite-responsive transcription factors (TF) that control expression of nutrient uptake enzymes in *Escherichia coli*, taken from EcoCyc [1]. All these systems follow the schematic in Fig. 1A.

| TF | Name | TF autoregulation | Operon inhibited by TF | Sequestering metabolite |
|---|---|---|---|---|
| ArsR | Arsenate inducibility regulator | Negative | arsB, from arsRBC operon | Arsenite / Antimonite ion |
| AlsR | Allose utilization regulator | Negative | alsABC, from alsRBACE operon | D-allose |
| BetI | Betaine Inhibitor | Negative | betT | Choline |
| ChbR | Chitobiose regulator | Negative | chbBCA from chbBCARFG operon | N,N'-diacetylchitobiose 6-phosphate |
| CytR | Cytidine regulator | Negative | nupC and nupG | Cytidine |
| FadR | Fatty acid degradation regulon | Negative | fadD | Acyl-CoA |
| GntR | Gluconate repressor | None, constitutive | gntT, gntU | D-Gluconate |
| LacI | Lactose inhibitor | None, constitutive | lacZYA | Allolactose |
| LldR | Lactate regulator | Negative | lldP, from lldPRD operon | S-lactate |
| LsrR | Quorum sensing system | Negative | lsrACDB operon | AI-2 (autoinducer) |
| NagC | N-acetylglucosamine transcriptional regulator | Negative | chbF, from chbBCARFG operon | Acetyl-D-glucosamine 6-phosphate |
| NanR | N-acetyl-neuraminic acid regulator | None, constitutive | nanT, from nanATEK-yhcH operon | N-acetylneuraminate |
| PaaX | Phenylacetic acid regulator | Negative | paaK, from paa operon | Phenylacetyl-CoA |
| PuuR | Putrescine utilization and transport regulator | Negative | puuP, from puuAP operon | Putrescine |
| RbsR | Ribose repressor | Negative | rbsACB, from rbs operon | D-ribose |
| SrlR | Glucitol Repressor | Negative | srlAEB, from srlAEBD-gutM-slrR-gutQ operon | D-sorbitol |
| TreR | Trehalose repressor | None, constitutive | treB, from treBC operon | Trehalose 6-phosphate |
| UlaR | Utilization of l-ascorbic acid repressor | None, constitutive | ulaABC, from ulaABCDEF operon | L-ascorbate 6-phosphate |

### S2. Kinetic model of fatty acid uptake.

To model the system in Fig. 1(B), we use the kinetic model:

$$\frac{dR}{dt} = P_R(R, p_r) - S - \mu \cdot R, \tag{E1}$$

$$\frac{dD}{dt} = b_D + \frac{a_D}{1+(K_D \cdot R)^{n_D}} - \mu \cdot D, \tag{E2}$$

$$\frac{dA}{dt} = \frac{k_{cat,D} \cdot \mathrm{OA}}{K_{m,D}+\mathrm{OA}} \cdot D - \frac{k_{cat,B} \cdot A}{K_{m,B}+A} \cdot B - 2 \cdot S - \mu \cdot A, \tag{E3}$$

$$\frac{d\mathrm{sR}}{dt} = S - \mu \cdot \mathrm{sR}, \tag{E4}$$

$$S = k_f \cdot R \cdot A^2 - k_r \cdot \mathrm{sR}, \tag{E5}$$

where *R*, *D*, *A* and aR represent the concentrations of transcription factor FadR, uptake enzyme FadD, internalized fatty acyl-CoA and sequestered acyl-CoA-FadR complex, respectively (Fig. 1B). During inducting, two molecules of acyl-CoA bind to sequester 1 dimer of FadR [2]. We model this reversible binding as mass-action kinetics (Eq. E5). The term $P_R(R, p_r)$ represents the expression and autoregulation of the fadR promoter. To model TF expression when under negative autoregulation (*n*), positive autoregulation (*p*) or constitutive expression (*c*), we write

$$P_{R,n} = b_n + \frac{a_n}{1+(K_n \cdot R)^{n_R}}, \tag{E6}$$

$$P_{R,p} = b_p + \frac{a_p \cdot (K_p \cdot R)^{n_R}}{1+(K_p \cdot R)^{n_R}}, \tag{E7}$$

$$P_{R,c} = p_c, \tag{E8}$$

respectively. We can use the model to simulate growth in continuous culture by fixing oleic acid concentration (OA) in Eq. E3. Model parameters can be found in Table S2.

**Table S2. Model parameters.**

| **Parameter** | $b_R$ | $a_R$ | $K_R$ | $n_R$ |
|---|---|---|---|---|
| **Description** | fadR basal exp. rate<br>Units = $\mu M/h$<br>*(Fitted)* | fadR promoter strength<br>Units = $\mu M/h$<br>*(Fitted)* | Affinity of FadR for its own promoter<br>Units = $1/\mu M$<br>*(Fitted)* | Hill coefficient<br>Units = N/A<br>*(Fixed = 1)* |
| **Parameter** | $b_D$ | $a_D$ | $K_D$ | $n_D$ |
| **Description** | fadD basal exp. rate<br>Units = $\mu M/h$<br>*(Fitted)* | fadD promoter strength<br>Units = $\mu M/h$<br>*(Fitted)* | Affinity of FadR for fadD promoter<br>Units = $1/\mu M$<br>*(Fitted)* | Hill coefficient<br>Units = N/A<br>*(Fixed = 2)* |

| Parameter | $k_{cat,D}$ | $K_{m,D}$ | $k_{cat,B}$ | $K_{m,B}$ |
|---|---|---|---|---|
| **Description** | Turnover rate of FadD<br>Units = $1/h$<br>*(Fitted)* | Michaelis const. for FadD<br>Units = $\mu M$<br>*(Fitted)* | Turnover rate of PlsB enzyme<br>Units = $1/h$<br>*(Fitted)* | Michaelis const.<br>Units = $\mu M$<br>*(Fitted)* |
| **Parameter** | $B$ | $k_f$ | $k_r$ | $\mu$ |
| **Description** | Conc. of PlsB enzyme<br>Units = $\mu M$<br>*(Fixed = 0.1369)* | Fwd rate of sequestering<br>Units = $1/h$<br>*(Fitted)* | Reverse rate of sequestering<br>Units = $1/h$<br>*(Fitted)* | Cell growth rate<br>Units = $h^{-1}$<br>*(Fitted)* |

To fit model parameters, we use time course data from batch cultures induced with titrations of oleic acid, shown in Fig. S1(B). We used a red fluorescent protein (RFP) gene placed at 3' of the *fadD* promoter on a low copy number plasmid (pSfadDk-RFP). The plasmid was incorporated to a *fadE* knockout strain to make *ΔfadE*-reporter. The *fadE* knockout strain was chosen to reduce the consumption rate of intracellular acyl-CoA and to simplify the metabolite dynamics for this parameterization purpose. Cells were cultivated in M9 glycerol (M9G) medium, in flasks, to exponential growth phase and induced with varying concentrations of oleic acid. Time course measurements of RFP fluorescence (Fig. S1(B)) and cell density (Fig. S1(A)) were recorded.

For model fitting we extended the Eqs. (E1)-(E8) with population growth in batch culture:

$$\frac{dX}{dt} = \mu \cdot X, \quad \text{(E9)}$$

$$\frac{d\,\mathrm{OA}}{dt} = -\frac{k_{cat,D} \cdot \mathrm{OA}}{K_{m,D} + \mathrm{OA}} \cdot D \cdot X, \quad \text{(E10)}$$

with parameters defined in Table S2. We first converted fluorescence values to units of concentration (µM) by assuming that the average fluorescence value in the absence of inducer (dark blue points, Fig. S1(B)) represents the steady state concentration of FadD reported in [3], measured in cells grown in the same media as ours (M9G). This gives a conversion factor of $1.12 \times 10^{-4}$ µM per unit of fluorescence, which was then applied to all fluorescence values; results are in Fig. S1(B).

We then performed a weighted least-squares fitting of simulations to the data. We define $D_i(t, p)$ and $\bar{d}_i(t)$ as the simulated and average measured FadD concentration (from three biological replicates), at time t, from the $i^{th}$ time series. The index *i* = 1, … ,9 refers to the time course when induced with oleic acid = 0, 0.4, 1, 4, 10, 40, 100, 400, and 1000 µM respectively. Fitting was performed to find optimal values of model parameters (*p*) that minimize the cost function

$$C = \sum_{i=1}^{9} \sum_{t=0}^{t_{\text{end}}} \left( \frac{\bar{d}_i(t)}{d_i^{\text{SEM}}(t)} \right) \cdot \left( \frac{D_i(t,p) - \bar{d}_i(t)}{\max_t(\bar{d}_i(t))} \right)^2, \tag{E11}$$

given constraints

$$\text{LB} \leq p \leq \text{UB}, \tag{E12}$$

where LB and UB are lower and upper bounds on the parameter search space. The term $d_{i,t}^{\text{SEM}}$ is the standard error measured from triplicate data. The term $\frac{\bar{d}_i(t)}{d_i^{\text{SEM}}(t)}$ in Eq. E11 ensures that the difference between simulation and data at each time point is weighted by the inverse of the relative standard error. This increases the weight of those contributions to the cost where data has a lower measured error. We optimized parameters with a two-step approach. We first used a genetic algorithm (GA) from the Global Optimization toolbox in MATLAB 2018a to find a candidate for a global minimum (using 200 generations of the GA), and then to initialize the solver `fmincon` and perform a local optimization using the same cost function (Eq. E11). Fitting was performed independently 100 times; results are shown in Fig. S1(B), and summary statistics of parameter values are given in Table S3.

Growth rates ($\mu$) were estimated through a least-squares fitting of the measured optical densities in Fig. S1(A) to the exponential function:

$$h(t) = h_0 \cdot e^{\mu t}. \tag{E13}$$

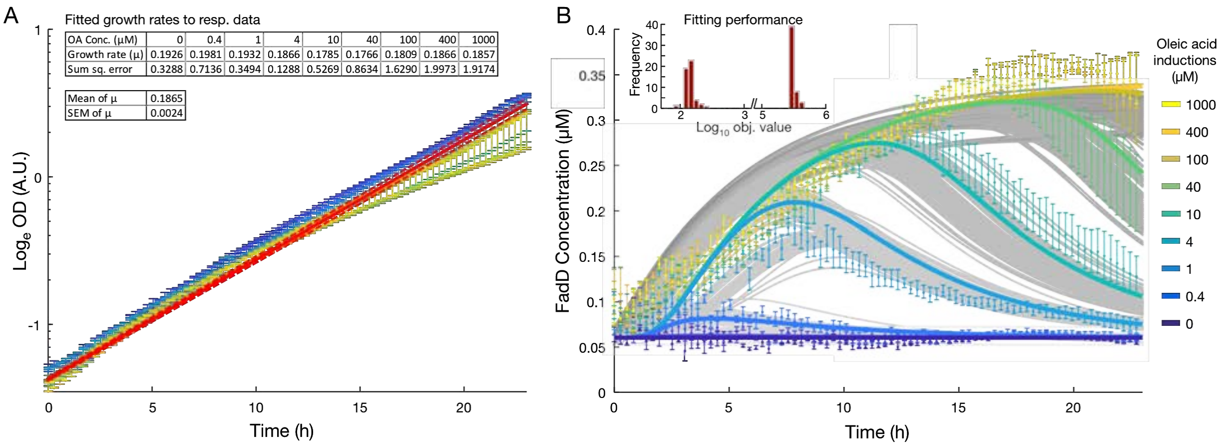


**Figure S1. Fitting model to data.** (A) Plot of time series data of measured optical density (OD), in log scale, during growth in media induced with titrations of oleic acid. Values of the fitted growth rate to each data series is given inset, including average growth rate (based on all data series) used to in the model; modelled growth shown in red line, ± SEM in red dashed lines. (B) Ensemble of 100 independent fits (gray curves) and the optimal fit (coloured curves) of simulations to time course data (point with errorbars), after converting fluorescence values to concentration. Error bars in data represent SEM from biological triplicates (n=3). Fitting performance in inset.

**Table S3. Results of parameter fitting.** Optimal parameter values together with search bounds and summary statistics for 100 independent fits. The bounds on growth rate, $\mu$, are based on ± two times the SEM from data (shown in Fig. S1(A), inset). Hill coefficients are fixed to $n_R$=1 and $n_D$=2, based on the number of FadR binding sites on the fadR and fadD promoters. Concentration of PlsB is fixed to 0.1369 µM, as taken from [3].

| Parameters | Optimal | Bounds of GA | | Summary Statistics | | | |
|---|---|---|---|---|---|---|---|
| | | Lower bound | Upper bound | Average | Median | SD | CV |
| $\mu$ | 0.1818 | 0.1817 | 0.1913 | 0.1854 | 0.1840 | 0.0036 | 1.9172% |
| $b_R$ | 0.0007 | 1.00E-06 | 0.0600 | 0.0210 | 0.0173 | 0.0140 | 66.7954% |
| $a_R$ | 0.0131 | 1.00E-06 | 0.1500 | 0.0414 | 0.0343 | 0.0343 | 82.8857% |
| $K_R$ | 4.3222 | 1.00E-03 | 100.0000 | 27.1436 | 22.9710 | 21.3182 | 78.5385% |
| $n_R$ | 1.0000 | - | - | - | - | - | - |
| $b_D$ | 0.0108 | 1.00E-06 | 0.1000 | 0.0108 | 0.0112 | 0.0015 | 13.5842% |
| $a_D$ | 0.0517 | 1.00E-06 | 0.1000 | 0.0486 | 0.0484 | 0.0021 | 4.4003% |
| $K_D$ | 305.9500 | 1.00E-03 | 750.0000 | 267.2850 | 215.9050 | 201.6395 | 75.4399% |
| $n_D$ | 2.0000 | - | - | - | - | - | - |
| $k_{catD}$ | 49.0000 | 1.00E-06 | 27,000.0 | 12,364.4 | 12,817.5 | 7,610.1 | 61.5483% |
| $Km_D$ | 0.0672 | 1.00E-02 | 650.0000 | 173.6743 | 149.1950 | 128.8831 | 74.2096% |
| $k_{catB}$ | 192.9100 | 1.00E-06 | 620.0000 | 235.0742 | 214.5650 | 160.4516 | 68.2557% |
| $Km_B$ | 45,429.0 | 1.00E-02 | 50,000.0 | 29,547.0 | 31,923.5 | 12,371.9 | 41.8719% |
| *PlsB* | 0.1369 | - | - | - | - | - | - |
| $k_f$ | 612.5500 | 1.00E-06 | 625.0000 | 409.3018 | 446.6350 | 174.9435 | 42.7419% |
| $k_r$ | 900.7300 | 1.00E-06 | 3,200.0 | 844.9572 | 515.6600 | 928.0828 | 109.8378% |
| Init. Biomass | 0.1648 | 1.00E-02 | 2.0000 | 0.2308 | 0.1521 | 0.2567 | 111.2037% |
| Obj Value | 82.0500 | - | - | 160,209.7 | 150,475.6 | 161,591.8 | 100.8627% |

To understand the impact of each model parameter on the recovery time, we conducted global parameter sensitivity analysis; see Supplementary Information S8 for details.

### S3. Impact of the exposure time to nutrient

As seen in Fig. S2, simulations suggest that for small increases in exposure time to oleic acid cause a decrease in recovery time, but for longer times recovery time is increased again. Further analysis of the simulations indicates that for exposure times, acyl-CoA accumulates to higher levels. In the OFF-state larger pools of accumulated acyl-CoA take longer to consume, which causes delays in the release of free FadR in the OFF-state. We infer that this delays the recovery of FadD, increasing recovery time. We hypothesized that the bottleneck lies in the consumption rate of acyl-CoA, as it is limited by the effective $v_{\max}$ of the consuming enzyme kinetics (in our system $v_{\max} = k_{\text{cat,B}}$PlsB). To computationally test this hypothesis, we increased the concentration of consuming enzyme (PlsB) in the model and found a faster release of free FadR which, in turn, reduced recovery time. Since longer exposure times increase the level of FadR stored in complex (a-R), we also found a general decrease in recovery time.

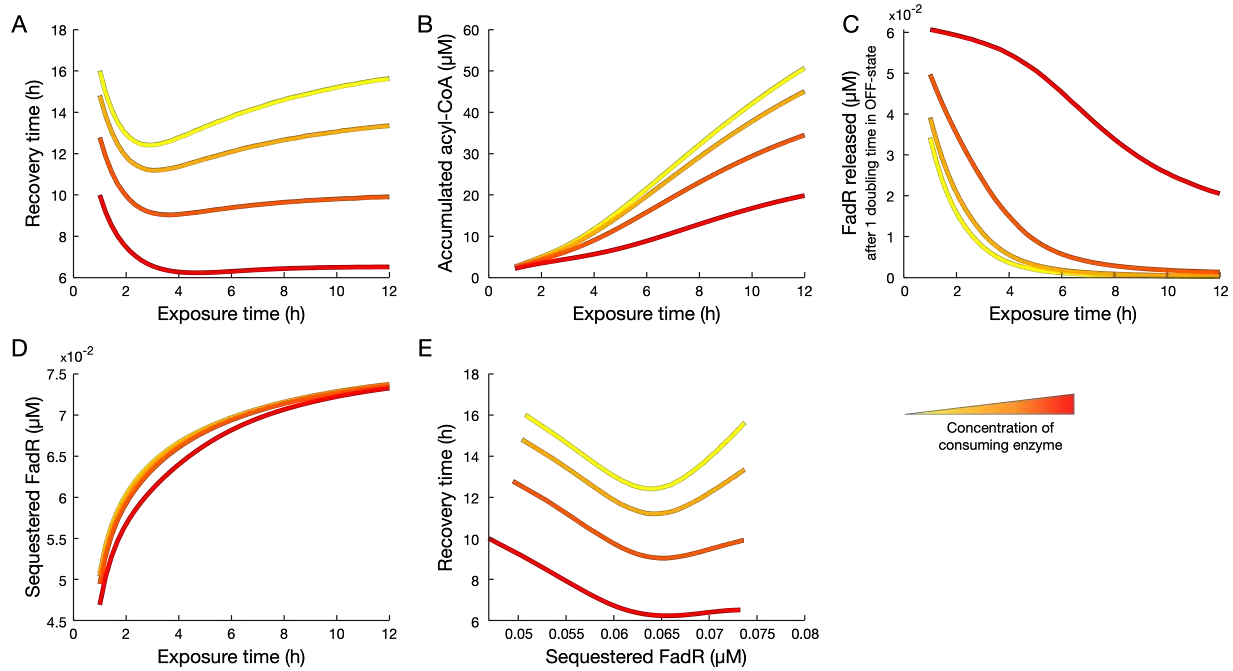


**Figure S2. Exposure time to oleic acid affects accumulated acyl-CoA and recovery time.** Coloured curves show how the respective relations are affected by increases in the concentration of consuming enzyme.

### S4. Plasmids and strains used in this study

#### Table S4. Plasmids used in this study.

| Plasmids | Replication Origin | Operon | Resistance | Reference |
|---|---|---|---|---|
| pSfadDk-RFP | SC101** | $P_{fadD}$-*rfp* | $Kan^R$ | This Study |
| pEfadRpoa-fadR | colE1 | $P_{fadRpo}$-*fadR* | $Amp^R$ | This Study |
| pSfadRpok-rfp | SC101** | $P_{fadRpo}$-*rfp* | $Kan^R$ | This Study |

#### Table S5. Strains used in this study.

| Strains | Relevant Genotype | Reference |
|---|---|---|
| *E. coli* DH1 | F- λ- *supE44 hsdR17 recA1 endA1 gyrA96 thi-1 relA1* | Hanahan 1983 |
| DH1(ΔfadE) | DH1, Δ*fadE* | Steen 2010 |

| | | |
|---|---|---|
| WT-reporter | DH1, pSfadDk-RFP | This Study |
| ΔfadE-reporter | DH1, Δ*fadE*, pSfadDk-RFP | This Study |
| PA-reporter | *DH1, fadR::$P_{fadRpo}$-fadR,* pSfadDk-RFP, pEfadRpoa-fadR | This Study |
| PA-FadR reporter | DH1, *fadR::$P_{fadRpo}$-fadR,* pSfadRpok-RFP, pEfadRpoa-fadR | This Study |

### S5. Steady state analysis: autoregulation affects FadR levels during induction

Here, we ask how the mode of FadR autoregulation affects how the pool of total FadR changes for long exposure times. To quantify this change, we look at the difference in the steady state $\Delta R_{T,ss} = R_{T,ss}^{(I)} - R_{T,ss}^{(0)}$ between the concentration of total FadR achieved in the ON-state ($R_{T,ss}^{(I)}$) to that achieved before induction ($R_{T,ss}^{(0)}$) for each of the three systems (negative autoregulation, positive autoregulation and constitutive expression).

To derive the expression for $\Delta R_{T,ss}$, we define total FadR as $R_{\mathrm{T}} = R + sR$, which from Eq. E1 and E4 follows

$$\frac{dR_{\mathrm{T}}}{dt} = p_R(R,p) - \mu \cdot R_{\mathrm{T}}, \quad \text{(E14)}$$

where $p_R(R,p)$ is the FadR synthesis rate as a function of free FadR and parameters, $p$, defined in Eqs. E6-E8 for each architecture. $\mu$ is growth rate and assumed constant before and during the ON-state. At steady state Eq. E14 gives

$$R_{T,ss} = \frac{p_R(R_{ss},p)}{\mu}. \quad \text{(E15)}$$

We can now write down an expression for $\Delta R_{T,ss}$ as

$$\Delta R_{T,ss} = R_{T,ss}^{(I)} - R_{T,ss}^{(0)} = \frac{p_R(R_{ss}^{(I)},p)}{\mu} - \frac{p_R(R_{ss}^{(0)},p)}{\mu}, \quad \text{(E16)}$$

where $R_{ss}^{(0)}$ and $R_{ss}^{(I)}$ are free FadR before induction and during the ON-state, respectively. In general, Eq. E1 at steady state gives

$$0 = p_R(R_{ss},p) - \mu \cdot R_{ss} - k_f \cdot A_{ss}^2 \cdot R_{ss} + k_r \cdot \mathrm{sR}_{ss}, \quad \text{(E17)}$$

where $R_{ss}$, $A_{ss}$ and $sR_{ss}$ are steady state concentrations of free FadR, acyl-CoA and sequestered FadR, respectively. Before induction, we have that $A_{ss} = 0$ and $\mathrm{sR}_{ss} = 0$ for the three architectures. Substitution into Eq. E17 leads to

$$0 = p_R(R_{ss}^{(0)},p) - \mu \cdot R_{ss}^{(0)}. \quad \text{(E18)}$$

We now substitute Eqs. E6-E8 into Eq. E18 for each mode of autoregulation, to get:

$$R_{ss,c}^{(0)} = \frac{p_c}{\mu} \text{ (constitutive)} \quad \text{(E19)}$$

$$R^{(0)}_{ss,n} = \frac{\left(\frac{K_n b_n}{\mu}-1\right)+\sqrt{\left(\frac{K_n b_n}{\mu}-1\right)^2+4\frac{K_n}{\mu}(a_n+b_n)}}{2K_n} \text{ (negative autoregulation)} \quad \text{(E20)}$$

$$R^{(0)}_{ss,p} = \frac{\left(\frac{K_a(a_a+b_a)}{\mu}-1\right)+\sqrt{\left(\frac{K_a(a_a+b_a)}{\mu}-1\right)^2+4\frac{K_a}{\mu}b_a}}{2K_a} \text{ (positive autoregulation)} \quad \text{(E21)}$$

To compute the steady state during the ON-state, we first solve for *sR* in steady state from Eq. E4 to obtain $\mathrm{sR}_{ss} = \frac{k_f A_{ss}^2 R_{ss}}{k_r+\mu}$, and then substitute into Eq. E17:

$$0 = p_R(R_{ss}, p) - \lambda(A_{ss}) \cdot R_{ss}, \quad \text{(E22)}$$

where we have defined

$$\lambda = \mu \cdot \left(1 + \frac{k_f A_{ss}^2}{k_r+\mu}\right), \quad \text{(E23)}$$

We now substitute Eqs. E6-E8 into Eq. E2 for each mode of autoregulation, to get the steady state concentration of $R^{(I)}_{ss}$ for each mode of autoregulation:

$$R^{(I)}_{ss,c} = \frac{p_c}{\lambda} \text{ (constitutive expression)} \quad \text{(E24)}$$

$$R^{(I)}_{ss,n} = \frac{\left(\frac{K_n b_n}{\lambda}-1\right)+\sqrt{\left(\frac{K_n b_n}{\lambda}-1\right)^2+4\frac{K_n}{\lambda}(a_n+b_n)}}{2K_n} \text{ (negative autoregulation)} \quad \text{(E25)}$$

$$R^{(I)}_{ss,p} = \frac{\left(\frac{K_a(a_a+b_a)}{\lambda}-1\right)+\sqrt{\left(\frac{K_a(a_a+b_a)}{\lambda}-1\right)^2+4\frac{K_a}{\lambda}b_a}}{2K_a} \text{ (positive autoregulation)} \quad \text{(E26)}$$

We now use the expressions for $R^{(I)}_{ss}$ in Eqs. E19-E21 and for $R^{(0)}_{ss}$ in Eqs. E24-E26 to compute the direction of change in steady state concentration of free FadR. From Eq. E23 we have that $\lambda \geq \mu$ for positive parameters, and therefore:

$$R^{(I)}_{ss,c} - R^{(0)}_{ss,c} \leq 0, \quad \text{(E27)}$$

$$R^{(I)}_{ss,n} - R^{(0)}_{ss,n} \leq 0, \quad \text{(E28)}$$

$$R^{(I)}_{ss,p} - R^{(0)}_{ss,p} \leq 0. \quad \text{(E29)}$$

Therefore, irrespective of the mode of autoregulation, the steady state level of free FadR is always lower in the ON-state relative to its steady state level before induction.

We can now return to Eq. E16 to compute the change in total FadR levels for each mode of autoregulation. **For constitutive expression**, substituting Eq. E8 into E16 we get

$$\Delta R_{T,ss,c} = \frac{p_c}{\mu} - \frac{p_c}{\mu} = 0, \quad \text{(E30)}$$

and therefore, there is no change in the level of total FadR in the ON-state, relative to its level before induction.

**For negative autoregulation**, substitution of Eq. E6 into E16 gives

$$\Delta R_{T,ss,n} = \frac{a_n}{\mu(1+K_n R_{ss,n}^{(I)})} - \frac{a_n}{\mu(1+K_n R_{ss,n}^{(0)})}. \quad \text{(E31)}$$

Using the relation in Eq. E28, it can be shown that $\Delta R_{T,ss,n} \geq 0$ and therefore the level of total FadR is increased in the ON-state, relative to its level before induction.

**For positive autoregulation**, substitution of Eq. E7 into E16 leads to

$$\Delta R_{T,ss,p} = \frac{a_p K_p R_{ss,p}^{(I)}}{\mu(1+K_p R_{ss,p}^{(I)})} - \frac{a_p K_p R_{ss,p}^{(0)}}{\mu(1+K_p R_{ss,p}^{(0)})}. \quad \text{(E32)}$$

Using the relation in Eq. E29, it can be shown that $\Delta R_{T,ss,p} \leq 0$ and therefore level of total FadR is decreased in the ON-state, relative to its level before induction.

In summary, we conclude that:

- For constitutive expression there is no change in total FadR and thus the system maintains the level of sequestered FadR during induction.
- For negative autoregulation, total FadR increases and thus the system builds up a larger pool of sequestered FadR during induction.
- For positive autoregulation, total FadR decreases and thus the system loses sequestered FadR during induction.

### S6. Construction and characterization of strain with positively autoregulated *fadR*

To engineer a strain with a self-activating architecture, a portion of the *fadR* promoter sequence, including the -10, -35 and the *fadR* operator sites were replaced with a sequence originating from the promoter of the *fabA* gene, which is positively regulated by FadR. To alter the genome sequence, we utilized pTarget-pCas genome editing system. The original promoter sequence and the engineered promoter ($P_{fadR\text{po}}$) sequence are show in Table S4. To enhance the expression of the FadR, we placed a plasmid copy of the positively regulated *fadR*, $P_{fadRpo}$-fadR, in a ColE1 origin plasmid. The positively autoregulated reporter strain (named "PA-reporter") was then created by transforming the pSfadDk-RFP reporter plasmid (Table S5, S6).

To confirm the self-activation of *fadR*, we measured the dose-response of the engineered $P_{fadRpo}$ promoter. A *rfp* gene with a strong ribosome binding site (RBS) was cloned to the 3' of the $P_{fadRpo}$ promoter in a BglBrick plasmid (pSfadRpok-rfp). The engineered promoter and RBS sequences are shown in Table S6. To measure dose-response output, PA-FadR reporter strain was grown in M9G with oleic acid concentrations = 0, 0.4, 1, 4, 10, 40, 100, 400 and

1000 µM. Cells were grown in a plate reader. Cell culture absorbance and RFP fluorescence were measured. Cultures were initially started at $OD_{600}$ = 0.001 and were allowed to reach steady state. Measurements for each culture condition were made in triplicate. Induction with high concentration of oleic acid reduces the output expression from the $P_{fadRpo}$ promoter (Fig. S3(A)), confirming $P_{fadRpo}$ as a positively autoregulated promoter.

**Table S6. Sequences of engineered promoter with positively autoregulated *fadR*.** (A) Native *fadR* promoter sequence, $P_{fadR}$. Bold lettering indicates FadR operator site, blue lettering indicates coding sequence. (B) Positively autoregulated *fadR* promoter, $P_{fadRpo}$, engineered in this work. The underlined sequence is derived from the *fabA* promoter region of *E. coli* DH1 genome. (C) Engineered $P_{fadRpo}$ used to control *rfp* expression.

| | |
|---|---|
| (A) | CCCTTTTTCTTCTTTTTGTCTGCTATCAGCGTAGTTAGCC**CTCTGGTATGATGAGTCC**AACTTTGTTTT GCTGTGTTATGGAAATCTCACTATGGTCATTAAGGCG |
| (B) | CCCTTTTTCTTCTTTTTATTCCG**AACTGATCGGACTTGTT**CAGCGTACACGTGTTAGCT ATCCTGCGTCAACTTTGTTTTGCTGTGTTATGGAAATCTCACTATGGTCATTAAGGCG |
| (C) | CCCTTTTTCTTCTTTTTATTCCG**AACTGATCGGACTTGTT**CAGCGTACACGTGTTAGCTATCCTGCGTCAACTTTGTTT TGCAGGTTTGTAAATAAAGGAGGGAGAAAGGGTATATGGCGAGTAGCGAA |

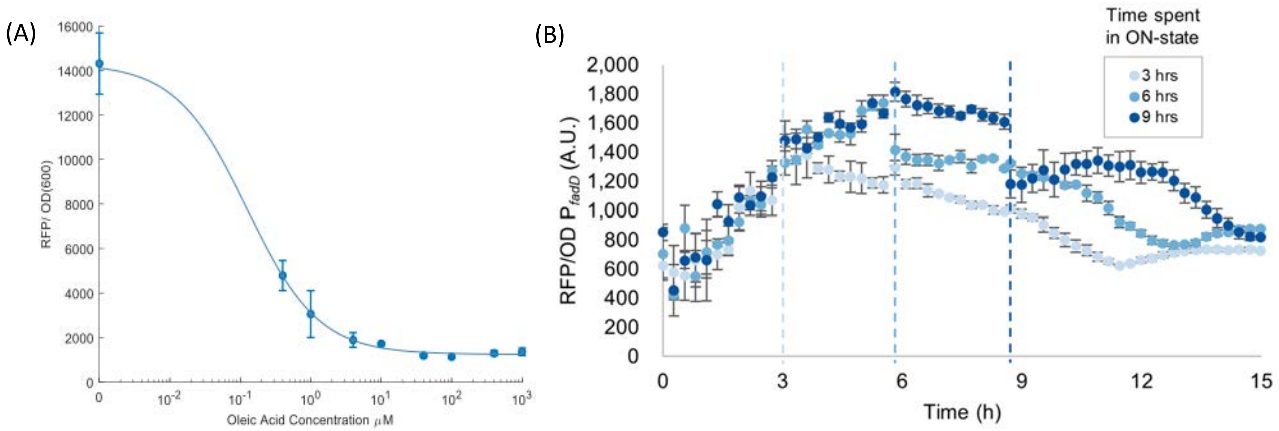


**Figure S3. Characterization and use of PA-reporter strain.** (A). Dose-response of $P_{fadRpo}$-rfp indicates FadR activated and OA inhibited $P_{fadRpo}$ expression. Error bars are SEM for biological replicates (n=3) and blue curve is fit of a hill equation to the mean oleic acid concentrations. (B). Time course fluorescence data for the switching experiment of the PA-reporter strain. Cells were induced by 1mM oleic acid at time zero, and grown for three different exposure times, 3, 6, and 9 hours (dashed vertical lines), after which cultures were rapidly switched to fresh media lacking oleic acid (OFF-state). Error bars represent the SEM from biological triplicates.

### S7. Steady state analysis: promoter strength affects level of sequestered FadR

For long exposure times, in Section S4 we have shown that constitutive expression and negative autoregulation can maintain or even build up the concentration of sequestered FadR. Here we study how the mode of autoregulation and parameters shape the steady state level of sR ($sR_{ss}$) achieved for long exposure times. We recall from the previous section that the steady state of sequestered FadR in the ON-state is

$$\mathrm{sR}_{ss} = \frac{k_f \cdot R_{ss} \cdot A_{ss}^2}{k_r + \mu}, \tag{E33}$$

where $R_{ss}$ is given by the formulae in Eq. E24 and E25 for constitutive and negative autoregulation

$$R_{ss,c} = \frac{p_c}{\mu \cdot \left(1 + \frac{k_f A_{ss,c}^2}{k_r + \mu}\right)}, \tag{E34}$$

$$R_{ss,n} = \frac{\left(\frac{K_n b_n}{\mu \cdot \left(1 + \frac{k_f A_{ss,n}^2}{k_r + \mu}\right)} - 1\right) + \sqrt{\left(\frac{K_n b_n}{\mu \cdot \left(1 + \frac{k_f A_{ss,n}^2}{k_r + \mu}\right)} - 1\right)^2 + 4 \frac{K_n}{\mu \cdot \left(1 + \frac{k_f A_{ss,n}^2}{k_r + \mu}\right)} (a_n + b_n)}}{2K_n}, \tag{E35}$$

respectively. Note that in E34-E35 we have substituted the expression for $\lambda$ given by Eq. E23. Substituting both expressions back into Eq. E33 we get

$$\mathrm{sR}_{ss,c} = \frac{p_c}{\mu} \cdot \frac{k_f A_{ss,c}^2}{k_f A_{ss,c}^2 + k_r + \mu}, \tag{E36}$$

$$\mathrm{sR}_{ss,n} = \frac{k_f}{k_r + \mu} \cdot \frac{1}{2K_n} \cdot \left( \left( \frac{K_n b_n}{\mu \left( \frac{1}{A_{ss,n}^2} + \frac{k_f}{k_r + \mu} \right)} - A_{ss,n}^2 \right) + \sqrt{ \left( \frac{K_n b_n}{\mu \left( \frac{1}{A_{ss,n}^2} + \frac{k_f}{k_r + \mu} \right)} - A_{ss,n}^2 \right)^2 + \frac{4 A_{ss,n}^2 K_n (a_n + b_n)}{\mu \left( \frac{1}{A_{ss,n}^2} + \frac{k_f}{k_r + \mu} \right)} } \right), \tag{E37}$$

for constitutive expression and negative autoregulation, respectively. To explore how parameters affect $sR_{ss}$ for long exposure time, we consider the scenario when there is a large accumulation of acyl-CoA from high levels of inducer in the media. We therefore approximate $sR_{ss}$ for each system by evaluating Eq. E36 and E37 in the limit $A_{ss} \to \infty$. For constitutive expression we get

$$\lim_{A_{ss,c} \to \infty} \mathrm{sR}_{ss,c} = \frac{p_c}{\mu}. \tag{E38}$$

For negative autoregulation, we further assume that the basal expression of *fadR* promoter is negligible ($b_n = 0$), so that Eq. E37 simplifies to

$$\mathrm{sR}_{ss,n} = \frac{k_f}{k_r + \mu} \cdot \frac{1}{2K_n} \cdot \left( -A_{ss,n}^2 + \sqrt{ A_{ss,n}^4 + \frac{4 A_{ss,n}^2 K_n a_n}{\mu \left( \frac{1}{A_{ss,n}^2} + \frac{k_f}{k_r + \mu} \right)} } \right), \tag{E39}$$

and thus we obtain

$$\lim_{A_{ss,n} \to \infty} \mathrm{sR}_{ss,n} = \frac{a_n}{\mu}. \tag{E40}$$

In summary, from Eq. E38 and E40 we conclude that in both architectures, sequestered FadR is scaled by the fadR promoter strength.

### S8. Sensitivity analysis of kinetic model

We used global sensitivity analysis (GSA) to quantify the impact of model parameters on the recovery time. We performed GSA using the method of extended Fourier amplitude sensitivity test (eFAST) [4, 5]. We adapted the MATLAB code reported in [5] and implemented eFAST to calculate the first-order and total-order sensitivity indices (Fig. S7). We assumed that each parameter could vary from 0.1- to 10-fold its fitted value, and model output was defined as the recovery time of FadD for the given input parameters. To enable us to determine which parameters the recovery time was statistically and significantly more sensitive to, we adopted the method of adding a dummy parameter to the model (details in [5]). This was used to compare to the sensitivities of other model parameters and a 2-tailed *t*-test was performed to test for significance. The parameters with significantly higher sensitivities are highlighted with an asterisk in Fig. S7. The results suggest that recovery time is more sensitive to parameters associated with the sequestering kinetics of FadR by acyl-CoA and the promoter strength of fadR promoter. It is also sensitive to the parameters representing the expression and regulation of FadD, but this is expected as it directly affects recovery time.

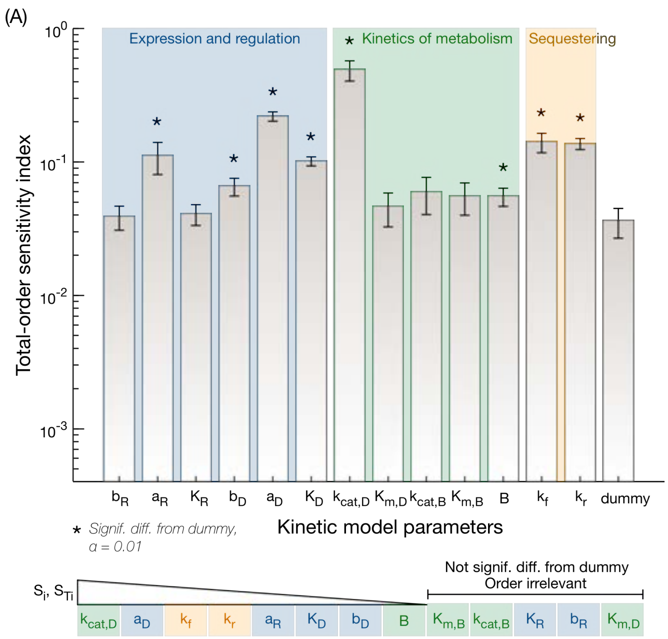


**Figure S4. Global sensitivity analysis of recovery time to model parameters.** Bar plot of the total-order sensitivity indices calculated from global sensitivity analysis (GSA) with eFAST. Sensitivities were calculated from 257 samples per search curve (set of parameters), and this sampling was repeated 7 times to ensure coverage of parameter values. Bars and error-bars show the average and 1 standard deviation of sensitivities over the 7 repeated sampling. eFAST assigns a dummy parameter a small, non-zero sensitivity (last bar). This was exploited to perform a two-tailed *t*-test to calculate whether the sensitivity of each parameter was significantly greater than that of the dummy parameter (asterisk, using a significance *α=0.01*). Parameters are listed in descending order of sensitivity (bottom).
